# Aqueous extract of *Phyllanthus niruri* protects against severe malaria by blocking erythrocyte invasion and modulating the host immune response

**DOI:** 10.1101/2021.07.09.451735

**Authors:** Jeje Temitope Olawale, Hironori Bando, Yasuhiro Fukuda, Ibukun Emmanuel Oluwafemi, Kentaro Kato

## Abstract

*Plasmodium falciparum* parasites are the major cause of malaria across Africa. Due to the appearance of multi-drug resistant parasites, new antimalarial drugs are needed. The medicinal plant *Phyllanthus niruri* is being used to treat fever and other symptoms of malaria in Nigeria; however, little is known about its antimalarial mechanisms. Here, we show that aqueous extract of *P. niruri* (PE) has multiple antimalarial effects, including anti-parasitic and host immunomodulatory activity. We found that co-culture of *P. falciparum* with PE drastically reduced parasite number, but PE did not inhibit parasite development or rupture; rather, it blocked erythrocytes invasion. In addition, we identified Astragalin as one of the antimalarial compounds which are contained in PE. Moreover, we found that PE suppresses the inflammatory activity and apoptosis of immune cells (T cells) and astrocytes and neurons in the central nervous system (CNS). Furthermore, we confirmed that oral administration of PE to mice suppressed parasite growth, excessive inflammation, CNS dysfunction, and the development of experimental cerebral malaria in an *in vivo* murine malaria model. Our findings demonstrate that PE has multiple effects on malaria progression, targeting both parasite and host cells.

## INTRODUCTION

Malaria is one of the most prevalent vector-borne infectious diseases; it is caused by the obligatory protozoan malaria parasite *Plasmodium*. The 2018 World Health Organization (WHO) report estimated that there were 405,000 deaths and 213 million cases of malaria around the world, and 90% of all malaria cases have occurred in sub-Saharan Africa^1^. Although human malaria is usually caused by four species of *Plasmodium* parasite—*P. falciparum, P. vivax, P. malariae*, and *P. ovale*—the majority of infections are caused by *P. falciparum*^2^. In addition, infection with *P. falciparum* can cause severe anemia, cerebral malaria (CM), and acute respiratory distress syndrome ^3–5^. Although there is currently no licensed malaria vaccine, some effective antimalarial drugs have been discovered, including quinine, which was extracted from the Cinchona tree^6^ and artemisinin, which was discovered in the Chinese plant *Artemisia annua*^7^. Thus, antimalarial drugs from medicinal plants have contributed to malaria control. However, the appearance of drug-resistant parasites has posed serious problems in recent years^8^, prompting the WHO to recommend artemisinin-based combination therapies (ACTs) for the treatment of malaria. Although most antimalarial drugs target the parasite inside the erythrocyte, at least one study suggests that an attractive target for therapeutic development is the step of merozoite erythrocyte invasion^9^, because inhibition of invasion would prolong exposure of the parasites to the immune system and drugs^10,11^. However, few antimalarial drugs target parasite erythrocyte invasion and new antimalarial therapeutics with targets that differ from those of existing antimalarial drugs are needed.

Historically, anti-pathogenic activity has been found in various medicinal plants^11^. In fact, some medicinal plants have been used for the treatment of malaria in endemic areas even though the science behind their efficacy is unclear^12,13^, suggesting that extracts from medicinal plants might be a promising source of new antimalarial compounds. *Phyllanthus niruri* L. (Euphorbiaceae) is a prevalent tropical plant^14^ with recognized medicinal properties^15^, being used for malaria treatment in some endemic areas. For example, in Nigeria, an aqueous extract obtained from the whole *P. niruri* plant has been used to prevent severe malaria or reduce malaria symptoms. Although, some previous studies have suggested that treatment with extract of *P. niruri* reduces the number of *P. falciparum in vitro* and murine *P. berghei* malaria *in vivo*^15,16^, the detailed mechanisms by which *P. niruri* inhibits parasite growth or suppresses the development of severe malaria are largely unknown.

*P. falciparum* infection induces dysfunction of the central nervous system (CNS), by, for example, disrupting the blood-brain-barrier (BBB) and causing neuronal damage in the brain, leading to CM^17^. *P. falciparum-associated* pathologies such as CM are the result of the complicated interaction between the host immune system and the parasite growth. Previous studies have shown that the development of CM is related to dysregulation of the immune system in a murine malaria model of experimental cerebral malaria (ECM)^18,19^. Mechanistic studies of ECM progression have reported that the expression of the inflammatory cytokine interferon-γ (IFN-γ) is critical for the development of ECM, because IFN-γ receptor-deficient mice are completely protected from ECM-dependent death^20,21^. In addition, it has been reported that T cell activation and production of IFN-γ are important for BBB disruption^22–24^. Thus, dysregulation of BBB function could cause the flow of cytokines, immune cells, and other toxic materials into the brain parenchyma, inducing CNS inflammation^25^. Therefore, an immunomodulatory approach to the suppression of *Plasmodium-induced* inflammation might be useful for malaria therapy. Moreover, a recent study suggested that the host cell is an attractive antimalarial drug target that could potentially prevent the acquisition of drug resistance by parasites^26^. Thus, the discovery of new antimalarial drug candidates with not only anti-parasitic activity but also host immunomodulatory activity is highly desirable.

Here, we found that aqueous extract of *P. niruri* (PE) can prevent the development of severe malaria by targeting both parasite and host cells. PE exerts anti-parasitic activity by blocking erythrocyte invasion by merozoites, and it read to inhibit CM progression. Importantly, both of these effects are driven by a different mechanism of action from that of most of the current antimalarial drugs, including artemisinin. Taken together, our data suggest that *P. niruri* might be useful in ACTs for malaria treatment and is a promising source for new antimalarial compounds.

## RESULTS

### PE treatment inhibits *P. falciparum* growth *in vitro*

The effect of *P. niruri* on *Plasmodium* parasite growth is largely unknown. Therefore, to test whether *P. niruri* has an anti-parasitic effect, freeze-dried aqueous *P. niruri* extracts (PE) were prepared (**Fig. 1A**). Previous reports indicated that PE does not show any toxicity *in vitro* or *in vivo*^27,28^. We confirmed that no significant cytotoxicity was observed at a concentration of 200 μg/ml PE against primary human foreskin fibroblasts (HFFs) (**Fig. 1B**). Next, to test the anti-parasitic activity of PE, *P. falciparum* were cultured with 200 μg/ml PE for 96 hours and then subjected to a growth inhibition assay (GIA) (**Fig. 1C**). As previously reported, PE treatment resulted in a significant reduction in the numbers of parasites compared with the nontreated condition (**Fig. 1C and Fig. S1A**). In addition, we found that the IC_50_ value of PE for *P. falciparum* inhibition was 35.11 μg/ml (**Fig. 1D**). A previous study reported that the extraction of *P. niruri* showed higher antimalarial activity (EC_50_: 2.9-4.1 μg/ml) compared to our result (EC_50_: 35.11 μg/ml)^16^. This disparity may be as a result of the difference in the materials and experimental procedure. For example, it is known that the metabolic profiles of *P. niruri* change with its geographical distribution^14^. Thus, we chose to use our materials and experimental procedure because our method and materials are suitable for detecting the effect and function of traditionally used *P. niruri* in Nigeria.

**Figure 1.**
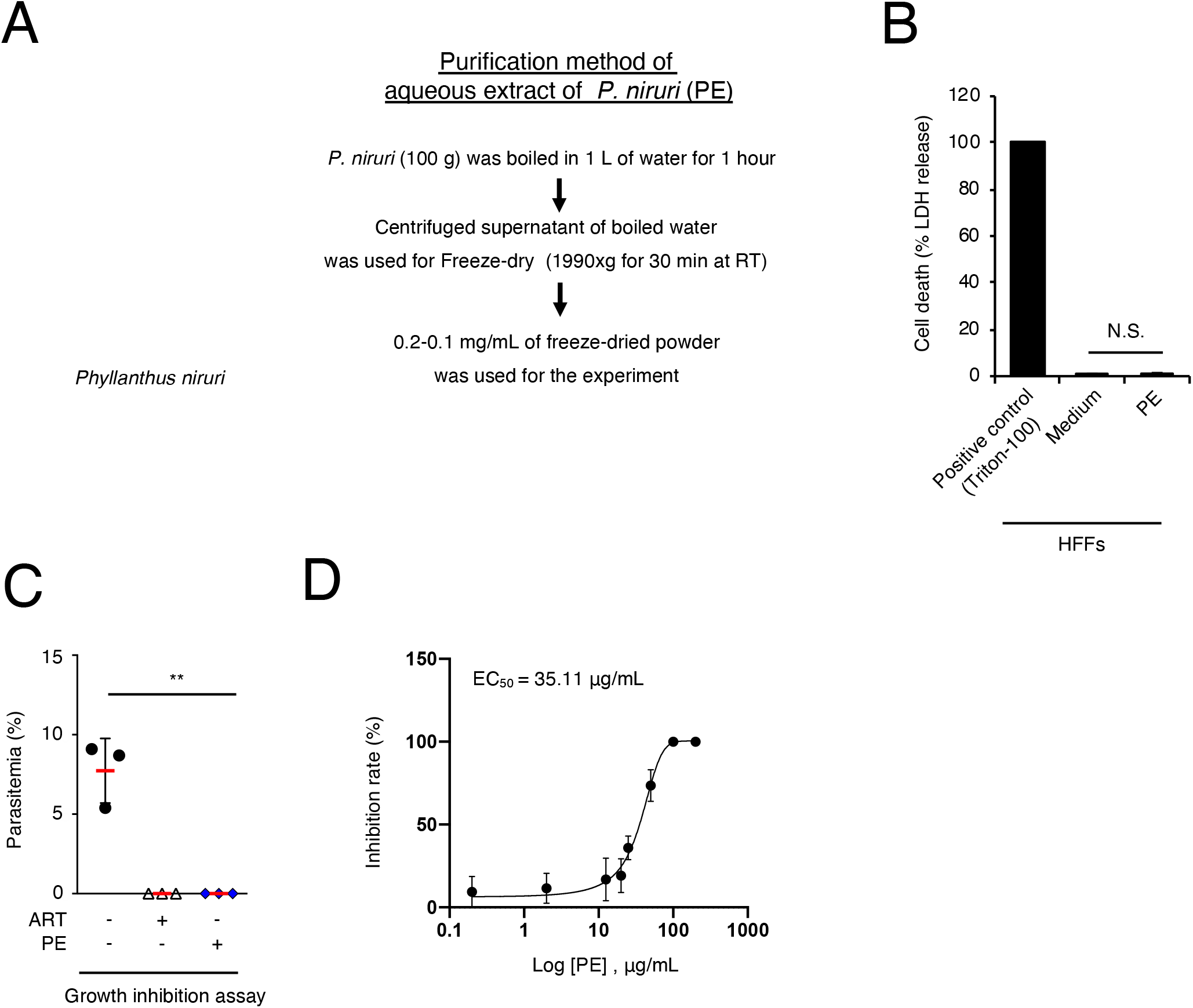

### PE affects erythrocyte invasion by *P. falciparum*

GIA is usually used to measure the comprehensive ability of a drug to inhibit parasite invasion, rupture, and/or development^29^. Therefore, to test whether the PE has an effect on parasite development inside the erythrocytes, synchronized ring formed parasites (immature trophozoites) were cultured with PE for 24 hours, and then compared the morphology of the parasites (**Fig. 2A**). There was no observable effect of the PE on the morphology of the parasites (**Fig. 2A**) or parasitemia (**Fig. S2A**), indicating that point of action of the PE is not parasite development into schizonts inside the erythrocytes. Next, we examined whether the PE has an effect on erythrocytes invasion by parasites. Synchronized schizonts were cultured with PE for 24 hours, and then parasitemia was assessed (**Fig. 2B**). PE treatment strongly reduced the parasite numbers (**Fig. 2B**). In addition, most of the infected parasites were not at the schizont stage but ring formed (**data not shown**), indicating that there was no effect of PE on parasite rupture or egress. Because erythrocyte invasion by parasites requires an interaction between host receptors and parasite ligands^30^, we assessed whether PE has an effect on erythrocyte surface receptors. The erythrocytes were treated with PE for 1 hour and then washed and cultured with infected erythrocytes for 24 hours (**Fig. 2C**). We found that pre-washed erythrocytes lost the ability to inhibit parasite invasion (**Fig. 2C**), suggesting that PE does not have an effect on erythrocyte surface receptors but on the parasite ligands or the parasite invasion processes.

**Figure 2.**
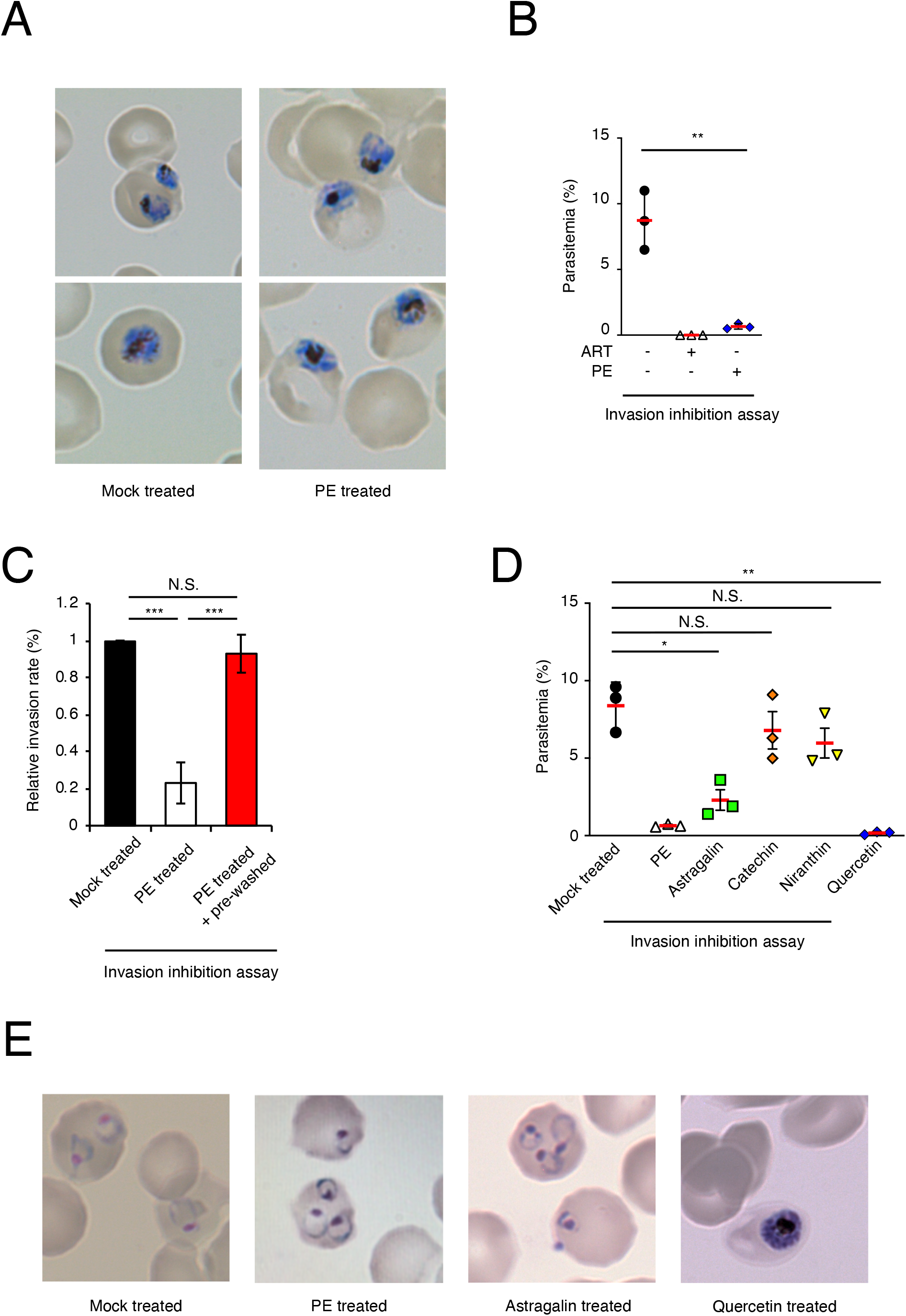

### Astragalin affects erythrocyte invasion by *P. falciparum*

Astragalin, Catechin, Niranthin and Quercetin have been reported to be abundantly identified in our study plant^14^. They are examples of polyphenolic compounds identified to be the constituent of the most active fraction of *P. niruri*^31^. We therefore subjected these compounds to invasion inhibition assay. We found that Astragalin and Quercetin, but not Catechin and Niranthin, had an antimalarial activity (**Fig. 2D**). Then we examined the antimalarial function of Astragalin and Quercetin. Astragalin showed an invasion inhibition activity with a mechanism similar to that of the aqueous extract of *P. niruri* (**Fig. 2E**). On the other hands, Quercetin showed its antiplasmodial effect by direct killing of the schizont, hence there was little or no egress (**Fig. 2E**). Taken together, we identified that Astragalin has an antimalarial activity in the similar way as PE. In addition, because we also found that *P. niruri* contains a number of antimalarial compounds, we focused on the antimalarial activity and function of whole plant (PE) extract to evaluate the effect and function of traditionally used *P. niruri* in natural.

### PE suppresses the activity of NF-κB dependent pro-inflammatory gene expression in T cells

We attempted to gain further insight into the mechanism by which PE prevents the development of severe malaria. Excessive gene or protein expression of pro-inflammatory cytokines, such as IFN-γ, has an important role in the dysregulation of the immune system and BBB destruction during *Plasmodium* infection^17,23^. Because T cells are an important source of IFN-γ to disrupt the BBB^32^, we examined the effect of PE on a human T-lymphoid cell line (MOLT-4). We first confirmed that PE treatment caused no significant cytotoxicity in MOLT-4 cells (**Fig. 3A**). Then, to assess whether PE treatment has an effect on IFN-γ gene expression, we cultured MOLT-4 cells with PMA and ionomycin for 3 hours (**Fig. 3B**). Although PMA/ionomycin co-stimulation showed marked induction of IFN-γ expression in PE non-treated MOLT-4 cells, PE treatment strongly suppressed IFN-γ induction (**Fig. 3B**), suggesting that PE plays an important role in the suppression of T cell inflammatory responses. NF-κB activation and its translocation from the cytoplasm to the nucleus is involved in IFN-γ expression^33^. Therefore, we co-stimulated PE-treated or non-treated MOLT-4 cells with PMA/ionomycin and compared the localization of NF-κB p65/RelA protein (**Fig. 3C**). The translocation of NF-κB p65/RelA protein to the nuclear compartment was observed in PE non-treated MOLT-4 cells that were co-stimulated with PMA/ionomycin. In sharp contrast, nuclear translocation of NF-κB p65/RelA was not observed in PE-treated MOLT-4 cells (**Fig. 3C**). To confirm the effect of PE on NF-κB activation, we generated a luciferase reporter plasmid harboring element dependent on NF-κB transcription factors. Then, we compared the luciferase activity induced by PMA/ionomycin co-stimulation in reporter plasmid-transfected MOLT-4 cells that were left untreated or were treated with PE (**Fig. 3D**). Although the luciferase activity in the PE nontreated MOLT-4 cells was increased by PMA/ionomycin stimulation, that in the PE-treated MOLT-4 cells was inhibited (**Fig. 3D**). In addition, there were no differences in the levels of NF-κB mRNA but significant differences in the levels of IκB-α, which is involved in the inhibition of NF-κB nuclear translocation^34^ (**Fig. S3A**). These results suggest that PE affects not the expression levels of NF-κB mRNA but the activation and/or nuclear translocation of NF-κB.

**Figure 3.**
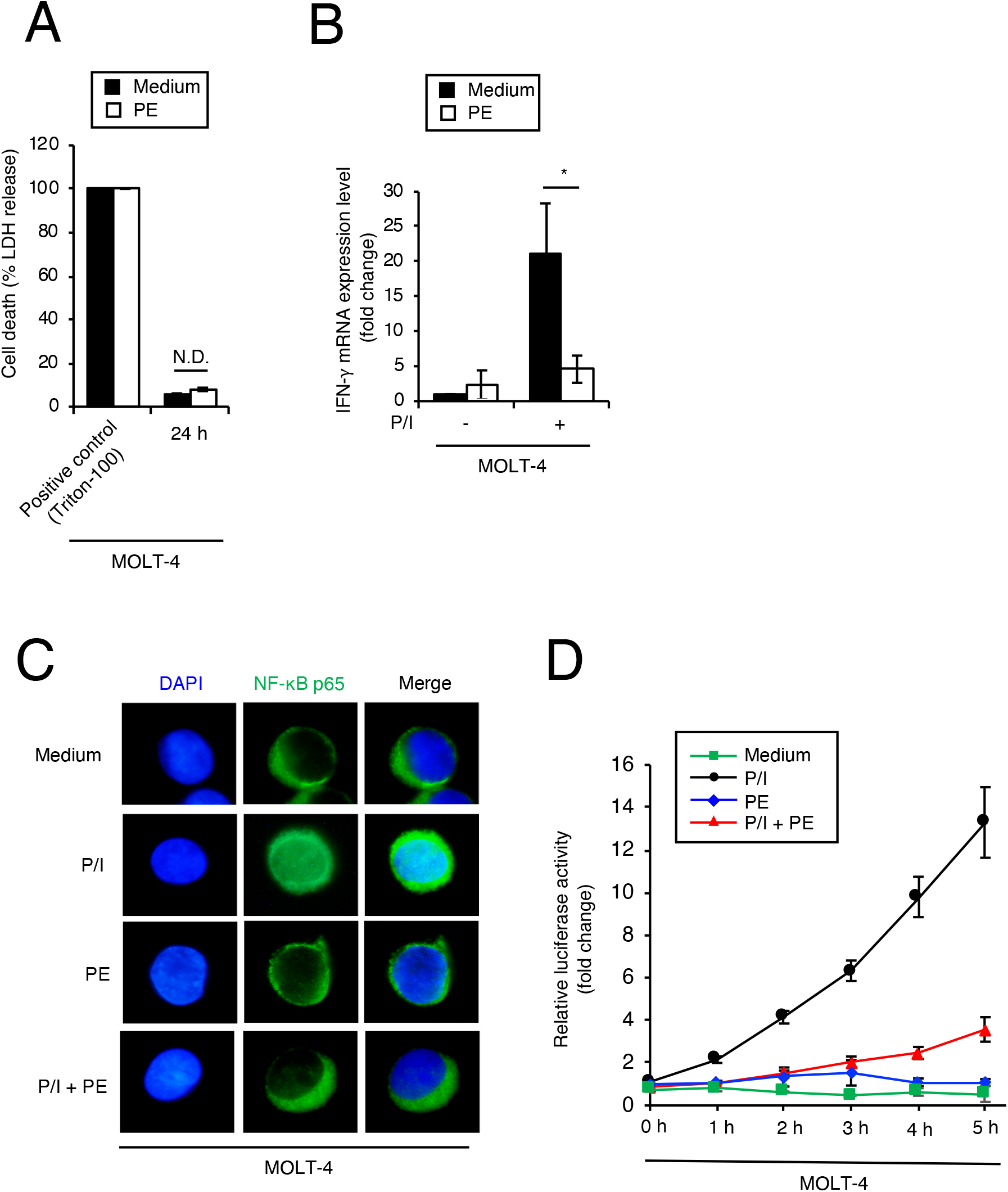

### PE blocks inflammatory activity and apoptosis in the CNS

The activation of astrocytes, a type of glial cell in the CNS, plays an important role in the progression of CM^35^. We confirmed that PE treatment caused no significant cytotoxicity in the human astrocyte cell line U-251 (**Fig. 4A**). Then, to assess whether PE treatment had an effect on IFN-γ gene expression, we cultured U-251 cells with PMA and ionomycin for 3 hours (**Fig. 4B**). Although co-stimulation with PMA/ionomycin markedly induced IFN-γ expression in PE non-treated U-251 cells, PE treatment inhibited IFN-γ induction (**Fig. 4B**), suggesting that PE also suppresses the inflammatory responses of astrocytes. CM progression induces neuronal apoptosis, leading to neuropathology^35,36^. We confirmed that PE treatment caused no significant cytotoxicity in the human neuron cell line IMR-32 (**Fig. 4C**). Then, to test whether PE could prevent neuronal apoptosis, we cultured IMR-32 cells with staurosporine with or without PE. Then, we compared the activity of caspase 3/7, which are proteases that initiate and execute apoptosis^37^ (**Fig. 4D**). We found that PE treatment severely reduced staurosporine-induced caspase 3/7 activity, indicating that PE inhibited the activation of the apoptotic pathway in neuron cells.

**Figure 4.**
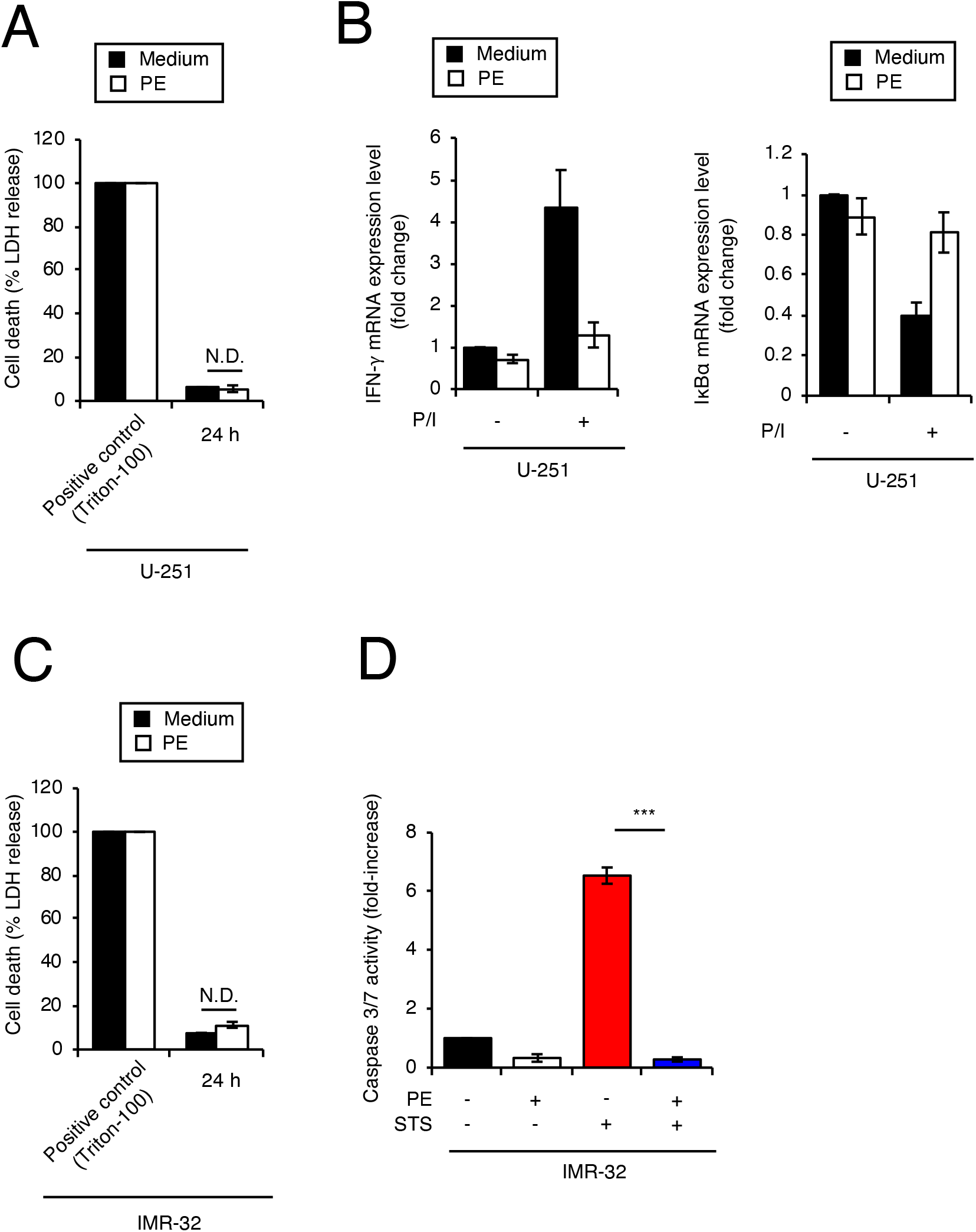

### PE has *in vivo* antimalarial effects against murine *P. berghei* malaria

Finally, we tested whether PE treatment could control the development of CM *in vivo*. At first, we assessed the immunomodulatory activity of PE by mice sepsis model because Sepsis progression has been linked to IFN-γ release, which is also responsible for systemic inflammatory syndrome through its ability to enlarge proinflammatory response^38^. We found out that the level of IFN-γ in the serum of PE treated group of mice was significantly reduced compared to the non-treated group (**Fig. S4A)**. Then, we assessed the antimalarial activity of PE by experimental cerebral malaria (ECM). The murine malaria parasite *P. berghei* ANKA was used to infect C57BL/6 (B6) mice, which are known to be a good model for ECM^19^. When Mock- or PE-treated mice were challenged with *P. berghei-infected* erythrocytes, the PE-treated mice exhibited reduced parasitemia (**Fig. 5A**) and higher survival rates (**Fig. 5B**). In addition, we found that there was significantly reduced disruption of the BBB in PE-treated mice (**Fig. 5C**). Furthermore, serum IFN-γ levels in *P. berghei*-infected mice were significantly reduced in the presence of PE (**Fig. 5D**). Taken together, these results indicate that PE induces multiple *in vivo* effects to prevent CM progression, including inhibition of parasite growth, suppression of excessive inflammation, and prevention of CNS dysfunction.

**Figure 5.**
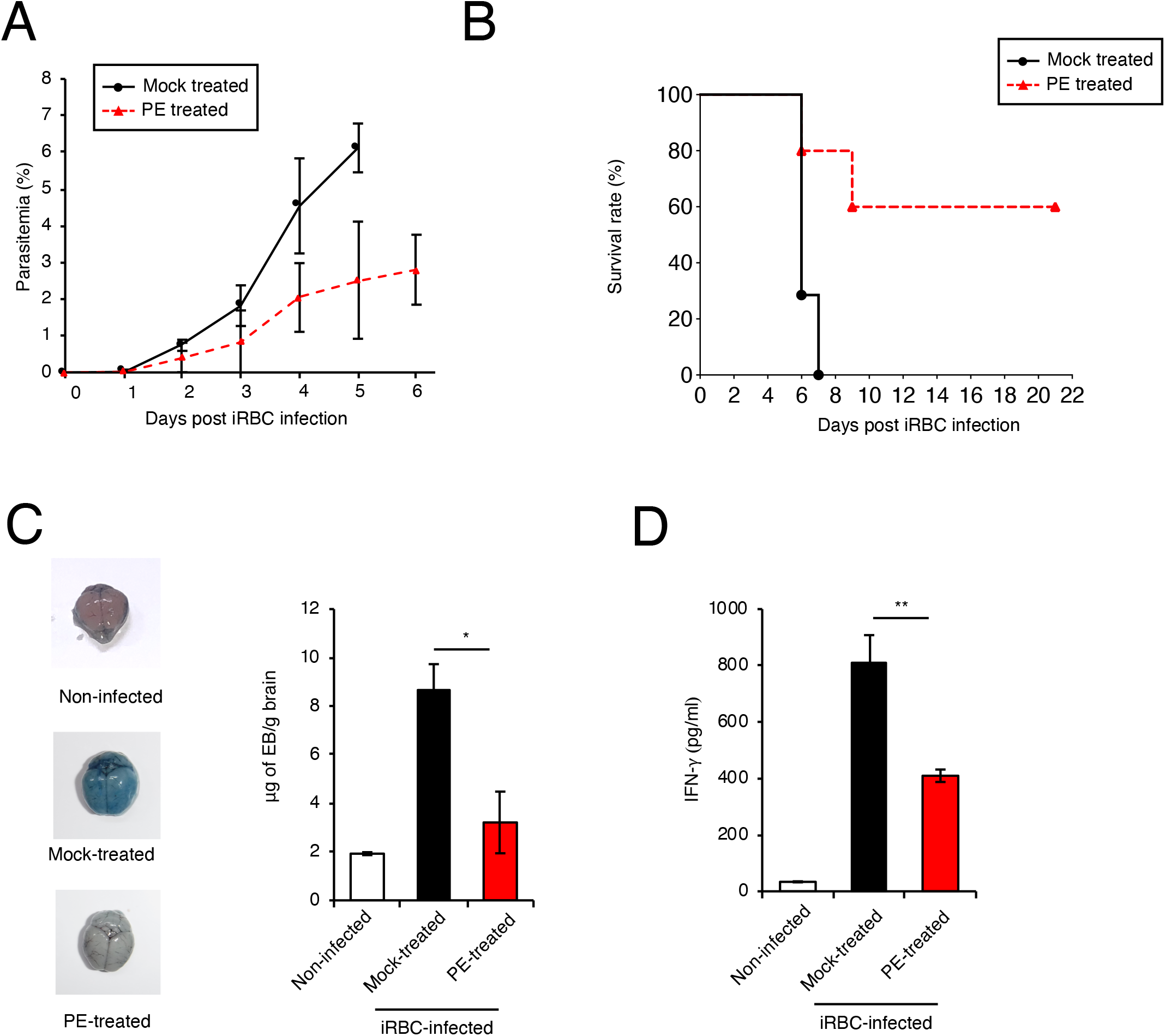

## DISCUSSION

The present study showed that an aqueous extract of *P. niruri* (PE) has multiple effects, both direct and indirect, against the disease progression pathway of severe malaria. PE has a direct effect in that it targets the parasite, by inhibiting erythrocyte invasion, and an indirect effect in that it targets the host via its immunomodulatory activity to suppress excessive inflammation and prevent CNS disfunction.

Over the years, antimalarial drugs against the blood stage of *P. falciparum* have been used for the treatment of malaria^29,39^. However, new antimalarial drugs are still required because of the emergence of multi-drug resistant parasites^9,40^. Accordingly, new candidate antimalarial drugs have been sought. Medicinal plants are a recognized repository of drug lead compounds against various diseases including malaria^41,42^. Scientific evaluation of these natural pharmaceuticals can offer an effective and low-cost route to obtain drugs for the treatment of diseases, including malaria^12^. One such medicinal plant, *Phyllanthus niruri*, is being used for antimalarial therapy in Nigeria. In this study, we confirmed the anti-parasitic activity of PE by using a *P. falciparum* GIA, which suggested that PE might have potential as an antimalarial drug.

The life cycle of the *Plasmodium* parasite in the host is separated into two different stages: the liver-stage and the blood-stage. The blood-stage is also known as the clinical stage^43^, in contrast to the liver stage, which is sometimes termed the silent stage^44^. *Plasmodium* merozoites repeat the cyclic process of erythrocyte invasion, development, and rupture, leading to lysis of the erythrocytes and severe anemia. Thus, although each step in the life cycle of the blood-stage parasite is attractive as an anti-parasitic target because it is an essential event for parasite infection, most of the existing antimalarial drugs target parasite development^9,45^. In contrast, little has been reported about compounds that target erythrocyte invasion by parasites. One report suggested that the chemical compound YA29 can affect erythrocyte invasion by merozoites;^46^ however, another study suggested that YA29 affects the rupture of schizonts^47^, and therefore, the mechanism of action of YA29 remains controversial. In the present study, we found that PE has an effect on erythrocyte invasion by parasites. Importantly, although the *Plasmodium* GIA is capable of detecting PE efficacy with respect to inhibition of invasion, development, and/or rupture^29^, it clearly showed that PE does not have an effect on the development and rupture of parasites. Furthermore, we found that PE did not have any effect on the erythrocyte cell surface receptors during the invasion process despite both the host receptor and parasite ligand being important for the erythrocyte invasion step^30^. This result is important for ensuring low cytotoxicity because inhibition of the function of erythrocyte cell surface receptors could lead to problems such as inducing hemolysis and prevention of the interaction with Fc receptors on phagocytic macrophages^48^. In addition, we found that Astragalin and Quercetin, which were found to be abundantly identified in this plant^31^, are active compounds against *Plasmodium* parasite. Because both compounds have never been reported as antimalarial compound, it is an important finding. Notably, we discovered that Astragalin is a novel active compound against *Plasmodium* with parasite invasion inhibition activity. This mechanism of antimalarial activity is quite an important finding because there is no such mechanism of action among the existing anti-malarial drugs and little is known about compounds which has such an activity. Taken together, these result strongly suggest that PE may have potential for use in combination with current antimalarial drugs, particularly ACTs because the antimalarial mechanisms of artemisinin is completely different^49^.

There are two major severe malaria symptoms caused by *P. falciparum* infection: severe malarial anemia and CM. Several malarial anemia involves multiple factors among which is direct disruption of erythrocytes by parasite growth^4^. In contrast, CM is an immune-mediated pathology, which is indirectly caused by the *P. falciparum* infection-dependent immune dysfunction^50^. Our study and some other previous ones have shown that PE directly affects *P. falciparum* growth^15,16^; however, the immunomodulatory role of PE during *Plasmodium* infection has been unclear. Some previous studies have reported that *P. falciparum* infection induces excessive secretion of the pro-inflammatory cytokine IFN-γ from immune cells or glial cells of the CNS^22,25,51^, thereby resulting in BBB disruption and CNS dysfunction. Thus, the prevention of severe malaria progression may hinge on how well the excessive inflammatory response is controlled^52^. In fact, immunomodulators that downregulate the hyper-activated immune system caused by parasite infection have had variable success^53^. Interestingly, PE has previously been reported to attenuate pro-inflammatory cytokine release from activated macrophages^54^, suggesting the possibility that PE might be involved in immune modulation during malaria infection. In the present study, we found that PE suppressed IFN-γ expression in both T cells (immune cells) and astrocytes (major glial cells in the CNS) and also *in vivo* LPS-induced mouse sepsis. Some phytochemicals, such as flavonoid compounds, have been reported as immunomodulators that downregulate pro-inflammatory cytokine expression through inactivation of NF-κB^55,56^. Our study showed that PE-dependent inhibition of IFN-γ gene expression is also mediated by preventing NF-κB activation, suggesting the possibility that PE contains flavonoid or flavonoid-like compounds. In addition to excessive inflammatory activity, CM leads to a severe neuropathology caused by neuronal apoptotic cell death, and, therefore, protection of neurons from apoptosis is also important for preventing severe malaria progression. In the present study, we showed that PE functions to protect neurons from apoptotic signals. Furthermore, we showed that PE has both anti-parasitic effects and immunomodulatory effects *in vivo* by using a murine malaria model. Notably, our *in vivo* results were obtained by using a simple method with real-life applicability: aqueous extraction of *P. niruri* and oral administration. *P. niruri* is widely distributed and easily obtained in endemic areas. Moreover, aqueous extraction of *P. niruri* and its oral administration is simple and low cost; therefore, these processes are feasible and readily applicable to most endemic areas. Thus, our study suggests that aqueous extraction of *P. niruri* would be useful for the integrated management of severe malaria.

In summary, here we demonstrated that PE can provide protection from the development of severe malaria via multiple effects, including inhibition of erythrocyte invasion by parasites and suppression of CNS dysfunction both *in vitro* and *in vivo*. PE can not only be readily applied in endemic areas as a medicinal plant against malaria, but it may also contribute to the development of future next-generation antimalarial drugs because it may contain one or more useful antimalarial compounds.

## MATERIALS AND METHODS

### Ethics statement

The animal experiments were approved by the Animal Research Committee of Tohoku University and conducted in accordance with the ARRIVE and specific institutional guidelines.

### Mice, cells, and parasites

Six-week-old female C57BL/6N mice were obtained from Charles River Laboratories Japan. All animal experiments were conducted with the approval of the Animal Research Committee of Tohoku University. MOLT-4 cells were maintained in RPMI (Nacalai Tesque) containing 10% heat-inactivated FBS, 100 U/ml penicillin, and 0.1 mg/ml streptomycin. U-251 cells were maintained in MEM (Invitrogen) containing 10% heat-inactivated FBS, 100 U/ml penicillin, and 0.1 mg/ml streptomycin, as previously described^57^. IMR-32 cells were maintained in MEM (Invitrogen) containing 10% heat-inactivated FBS, 1% NEAA (Invitrogen), 100 U/ml penicillin, and 0.1 mg/ml streptomycin, as previously described^57^. *P. berghei* ANKA and *P. falciparum* HB3 clone were obtained from the Malaria Research and Reference Reagent Resource Center (MR4; American Type Culture Collection, Manassas, VA). The parasites were cultured in medium containing RPMI 1640, 25 mM HEPES, 100 μM hypoxanthine, 12.5 μg/ml gentamicin sulfate supplemented with 5% (w/v) Albumax I, and 62.5 μg/ml of NaHCO_3_, as previously described^58^.

### Plant material and extract preparation

Fresh samples of whole *Phyllanthus* plants were collected from Ado-Ekiti, Nigeria. The whole plants were air-dried and then pulverized with a mortar and pestle.

### Reagents

Antibodies against NF-κB p65 (sc-8008) were obtained from Santa Cruz Biotechnology.

### GIA of *P. falciparum*

The antiplasmodial activity of the extract was assessed on cultured *P. falciparum* HB3 parasites by means of microscopy as previously described^58^. The extract was weighed, dissolved in distilled water, and filter-sterilized (0.45 μm Millipore filter) to make a stock solution with a concentration of 5000 μg/ml. The working solution (concentration, 200 μg/ml) was made by diluting the stock solution with complete parasite media. One hundred and fifty microliters of *P. falciparum* parasite culture suspension, earlier synchronized to the ring stage with 5% sorbitol, was aliquoted into the wells of a 96-well microtiter plate to a final hematocrit of 2% and parasitemia of 0.5% (with fresh Type AB erythrocytes). The cultures were incubated at 37°C, with 5% CO_2_ and 5% O_2_. At 48-hour post-incubation, 5 μl of complete medium was added to each culture. Artemisinin was used as a positive control whereas wells containing no drug but culture at the same parasitemia and hematocrit served as a negative control. Evaluation of the outcome of the *in vitro* GIA was carried out 96-hour post-incubation, when the cultures were mostly in the late trophozoite stage, by means of microscopy. The experiment was carried out 3 times and each experiment was done in triplicate.

### *P. falciparum* invasion assay

For the invasion assays, purified schizont-infected erythrocytes were prepared by using the Percoll-Sorbitol method^58–60^. Briefly, the medium was removed from the parasite culture by centrifugation at 650 × *g* for 5 minutes at room temperature. The packed erythrocytes were then resuspended to 50% hematocrit and put on a gradient made from 70% and 40% Percoll-Sorbitol solution. The gradient was then centrifuged at 16,200 × *g* for 20 minutes at 20°C with no brake. Infected erythrocytes were collected from the 40/70 interface and washed twice with incomplete medium. A smear was made to assess the purity and estimate the number of cells by use of a hemocytometer. Purified schizonts were mixed with complete medium to obtain a hematocrit of 1%, and fresh AB erythrocytes were added for a total parasitemia of 2%. Cultures (150 μl) containing the compounds, Niranthin (0.01 ug/ul), Quercetin (0.03 ug/ul), Astragalin (0.03 ug/ul), Catechin (0.05 ug/ul) and PE (0.2 ug/ul) were transferred into 96-well plates. The compounds were dissolved based on their solubility in DMSO, and the final DMSO concentration was 0.1%. The cultures were then incubated for 24 hours at 37°C, with 5% CO_2_ and 5% O_2_. Three independent assays were done in triplicate. Culture supernatants were aspirated and the cell pellets were smeared and stained with Giemsa stain. Parasitemia was measured at the ring stage (appropriately 24 hours post-incubation).

### *P. falciparum* development assay

For the *P. falciparum* development assay, purified schizont-infected erythrocytes were prepared by using the Percoll-Sorbitol method, as discussed above. Fresh AB erythrocytes were incubated with the plant extract for 1 hour after which they were washed three times. Purified schizonts were mixed with complete medium to obtain a hematocrit of 1%, and the washed erythrocytes were added for a total parasitemia of 2%. The non-treated group and a group without washing after treatment served as control groups. Cultures (150 μl) were transferred into 96-well plates and the extract was added to a final concentration of appropriately 200 μg/ml. Cultures were incubated for 24 hours at 37°C, with 5% CO_2_ and 5% O_2_. Three independent assays were done in triplicate. Culture supernatants were aspirated and the cell pellets were smeared and stained with Giemsa stain. Parasitemia was measured at the ring stage (appropriately 24 hours post-incubation).

### Assessment of *P. berghei* parasitemia

Blood smears were obtained daily from all infected mice and stained with Giemsa stain. Parasitemia was assessed by counting at least 3×10^3^ erythrocytes.

### Assessment of vascular leakage

Vascular leakage was measured as previously described^26^. Mice were injected intravenously with 200 μl of 2% Evans Blue (Nacalai Tesque) at 5 days post-infection with 5×10^6^ *P. berghei*-infected erythrocytes. Mice were sacrificed 1-hour post-Evans Blue injection; brains were isolated, photographed, and then weighed. Brains were then placed in 2 ml of formamide for 48 hours at 37°C. The Evans Blue concentration was measured at 620 nm in an Multi Detection Microplate Reader (Power scan HT, DS Pharma Biomedical) by using 100 μl of solution. The concentration of Evans Blue was calculated by using a standard curve starting at 200 μg/ml. Evans Blue content is expressed as “μg Evans Blue/g brain” in Figure 5D.

### Quantitative determination of cell viability (LDH assay)

Cytotoxicity induced by PE was measured as previously described^61^. Cell viability was monitored by measuring lactate dehydrogenase (LDH) leakage into the culture medium by using the CytoTox96 Non-Radio Cytotoxicity Assay kit (Promega). For adherent cells (HFFs, IMR-32 cells, and U-251 cells.), the cells were seeded in the plate at 150 μl and incubated at 37°C, with 5% CO_2_ and 5% O_2_. After 24 hours, the old media were aspirated and replaced with new media containing 200 μg/ml of extract. The cells and the extracts were then incubated at 37°C, with 5% CO_2_ and 5% O_2_ for various hours, as indicated. For the suspension cells (i.e., MOLT-4 cells), the cells were seeded in the plate at the appropriate culture volume with the extract at a final concentration of 200 μg/ml and incubated at 37°C, with 5% CO_2_ and 5% O_2_. Culture supernatant was collected and centrifuged at 20,000 × *g* for 5 for 5 min, and then the LDH activity in the cell-free supernatant was measured according to the manufacturer’s instructions by using Multi Detection Microplate Reader (Power scan HT, DS Pharma Biomedical). Cell-free medium was used as a negative control. Culture supernatant from TritonX-100 (0.1%) treated cells (to kill all the cells) was used as a positive control.

### IC_50_ determination for PE

To calculate the IC_50_ of PE against *P. falciparum*, the GIA assay was performed using 12.5, 25, 50, 100, and 200 μg/ml of PE. The parasite numbers in the PE-treated cells divided by those in the non-treated cells are shown as the “inhibition rate” in Figure 1D.

### Quantitative RT-PCR

Total RNA was extracted from the cells with Direct-zol RNA MicroPrep (Zymo Research), and cDNA was synthesized by using Verso Reverse transcription (Thermo Fisher Scientific). Quantitative RT-PCR was performed by using the Go-Taq Real-Time PCR system (Promega), as previously described^60^. The values were normalized to the amount of glyceraldehyde 3-phosphate dehydrogenase (GAPDH). The primer sequences are listed in Table S1.

### Measurement of NF-κB promoter activity

Cells were stimulated with PMA/ionomycin and luciferase activities in cell lysates were measured as previously described^26^. To measure the luciferase activity, whole cells were collected 0, 1, 2, 3, 4, and 5 hours post-stimulation, then lysed with 100 μl of lysis buffer (Promega) and sonicated. After centrifugation at 20,000 × *g* at 4°C, the luciferase activity of the supernatant was measured by using the Dual Luciferase Reporter Assay System (Promega) and a GLOMAX 20/20 luminometer (Promega).

### Immunofluorescence assays

MOLT-4 cells were cultured at 1×10^5^ cells per well, with or without drug treatment after stimulation with PMA/ionomycin for various periods of time. The cells were then pre-fixed with ice-cold methanol for 1 hour at −20°C, after which they were fixed with paraformaldehyde (PFA, 1% in PBS buffer) at 4°C for 1 hour. Next, the cells were blocked with 1% BSA in PBS at room temperature for 1 hour. The cells were thereafter incubated with the primary anti-NF-κB antibody for 1 hour at room temperature, followed by incubation with secondary antibodies (Anti-mouse FITC) for 1 hour at room temperature in the dark. After washing the cells with PBS, cells were stained with DAPI for 5 minutes at room temperature in the dark and observed with fluorescence microscopy (BX-40, Olympus/Axiocam 305, Zeiss).

### Transfection of reporter plasmids

pGL4 (NF-κB-luc2p), which drives the transcription of the Photinus luciferase reporter gene, and the Renilla luciferase-expressing plasmid pRL-TK were co-transfected into cells by using lipofectamine 2000 (Invitrogen).

### Detection of Caspase 3/7 activity

IMR-32 cells were seeded in 96-well plates (3×10^4^ cells/well). After 24 hours, the medium was changed and the cells were treated with PE (200 μg/ml) or staurosporine (1 μM) as a positive control. After 6 hours of treatment, induction of apoptosis was determined by measurement of caspase-3/7 activity using the luminometric Caspase-Glo 3/7 assay (Promega) according to the manufacturer’s protocol and a microplate reader (Microplate Reader SH900), as previously described^61^.

### Mouse sepsis model

Eight-week-old female C57BL/6N mice were obtained from Charles River Laboratories Japan. Sepsis was induced by LPS (20mg/kg) injection by intraperitoneally (i.p.). Mice were pre-treated with PE or PBS for 2 days by oral treatment. Four hours after LPS injection, serum were collected for ELISA.

### Quantification of IFN-γ in mouse serum

A mouse IFN-γ ELISA kit (RayBiotech) was used according to the manufacturer’s instructions. Briefly, 100 μl of mouse serum and the standard were added to a 96-well microplate coated with anti-mouse IFN-γ (immobilized antibody). The plate was then incubated at room temperature for 2.5 hours with gentle shaking. After washing 4 times, 100 μl of prepared biotinylated anti-Mouse IFN-γ was added and incubated at room temperature for 1 hour with gentle shaking. The wells were again washed 4 times and 100 μl of HRP-conjugated Streptavidin solution was added and incubated with gentle shaking for 45 minutes. The plate was then washed and One-Step substrate reagent (3,3,5,5-tetramethylbenzidine) was added in the dark with gentle shaking. After 30 minutes of incubation, the reaction was stopped by adding 50 μl of 2N sulphuric acid, and the color intensity was measured at 450 nm by using the microplate reader. A curve was prepared, plotting the absorbance at 450 nm against the concentration of IFN-γ in the standard wells. By comparing the absorbance of the samples to this curve, the concentration of IFN-γ in the samples was determined.

### Statistical analysis

All statistical analyses were performed by using Prism 8 (GraphPad), as previously described^26,61^. All experimental points and n values represent an average of three biological replicates (three independent experiments). The statistical significance of differences in mean values was analyzed by using an unpaired two-tailed Student’s t-test. And the Mann-Whitney U test or Kruskal–Wallis test was used to analyze the differences between means with 95% confidence interval. *P* values less than 0.05 were considered to be statistically significant. The statistical significance of differences in survival times of mice between two groups was analyzed by using the Kaplan-Meier survival analysis Log rank test.

---

**Supplemental Figure 1.**
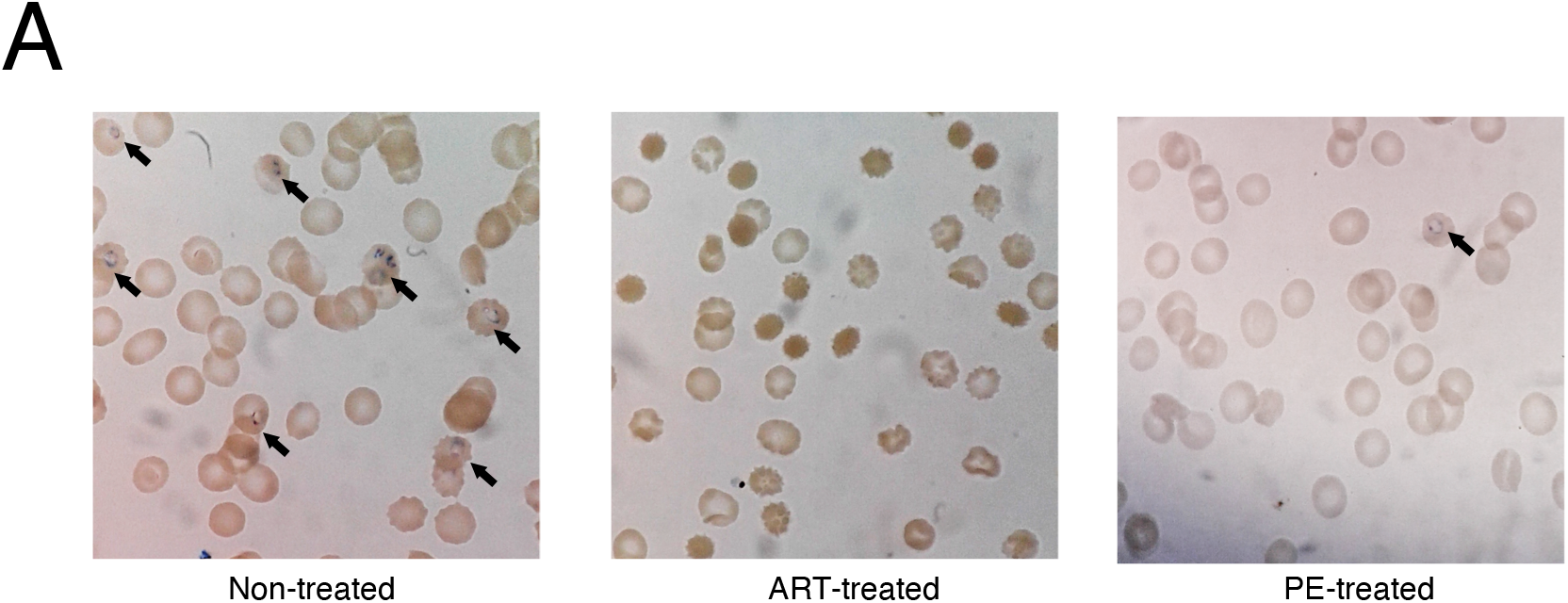

**Supplemental Figure 2.**
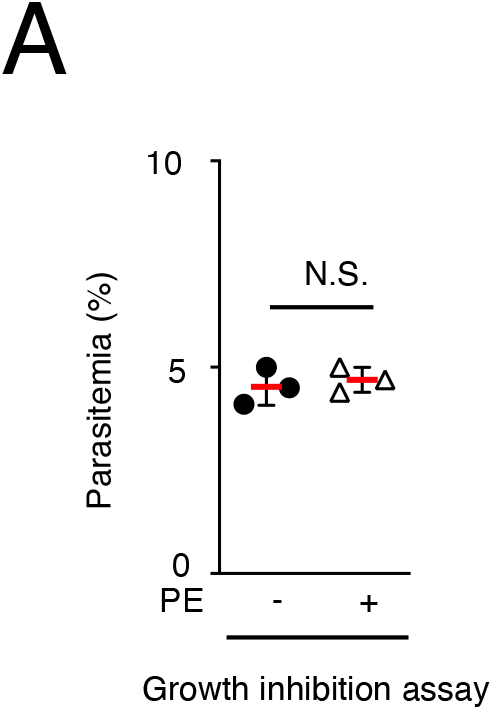

**Supplemental Figure 3.**
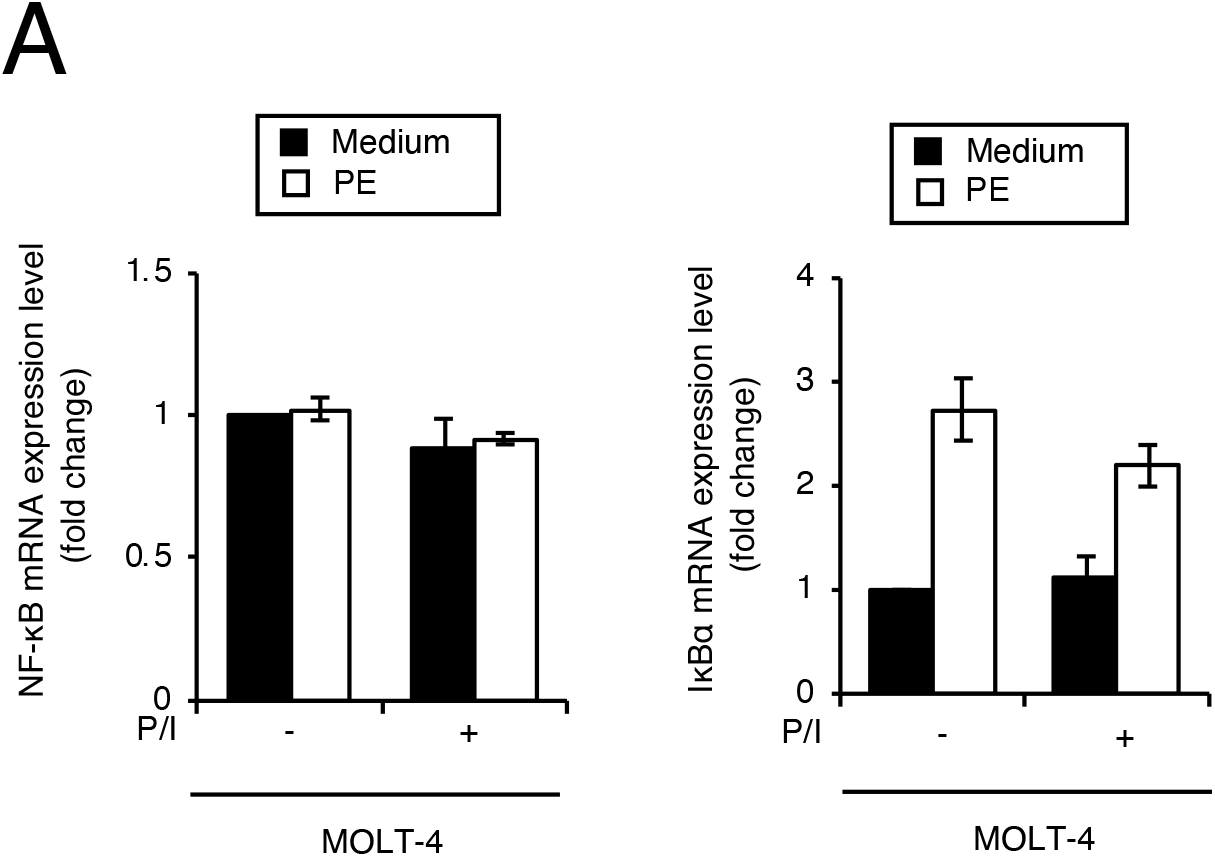

**Supplemental Figure 4.**
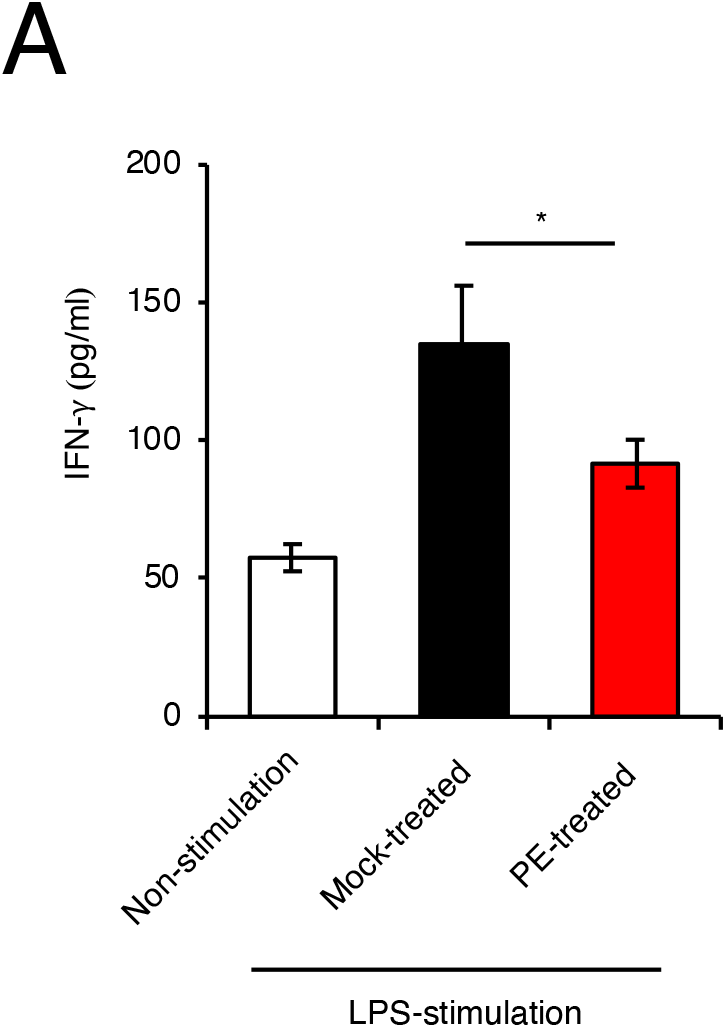

