## Supplementary material for "Aqueous extract of *Phyllanthus niruri* protects against severe malaria by blocking erythrocyte invasion and modulating the host immune response": supplental figure

P

**TITLE:**

**Authors' list:**

Temitope Olawale Jeje
Hironori Bando
Yasuhiro Fukuda
Emmanuel Oluwafemi Ibukun

Supplemental Figure 1_Jeje et al.

A

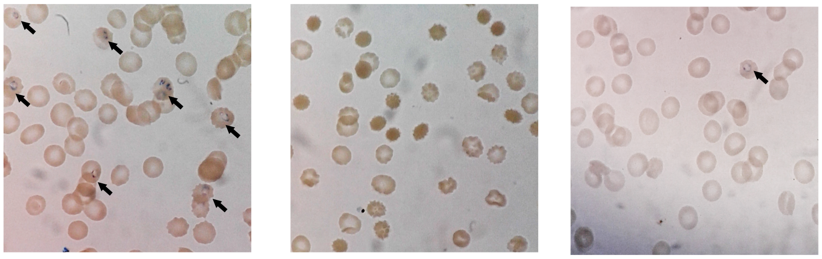

**Supplemental Figure 1. PE treatment inhibits *P. falciparum* growth *in vitro.***

Infected erythrocytes were treated or not with PE for 96 hours. Artemisinin (ART) was used as the positive control drug. The arrow indicates infected erythrocytes.

Supplemental Figure 2_Jeje et al.

A

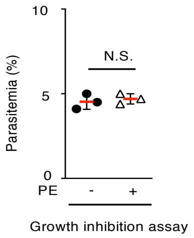

**Supplemental Figure 2. PE treatment does not affect the number of parasitemia *in vitro.***

Infected erythrocytes were treated or not with PE for 24 hours. There was no significant difference in parasitemia, as indicated. The indicated values are means ± s.d. (three biological replicates per group from three independent experiments). N.S., not significant; (Mann-Whitney U test).

### Supplemental Figure 3_Jeje et al.

A

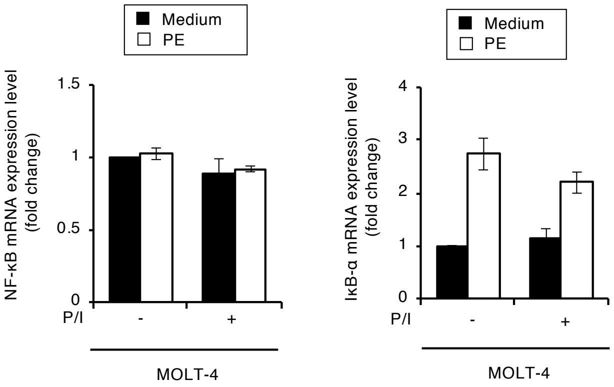

**Supplemental Figure 3. PE treatment affects the expression of IκB-α, but not that of NF-κB.**

Quantitative RT-PCR analysis of NF-κB and IκB-α mRNA levels in MOLT-4 cells that were untreated or treated with PE for 1 hour and then incubated for 3 hours with PMA and ionomycin (P/I). The indicated values are means ± s.d. (three biological replicates per group from three independent experiments).

### Supplemental Figure 4_Jeje et al.

A

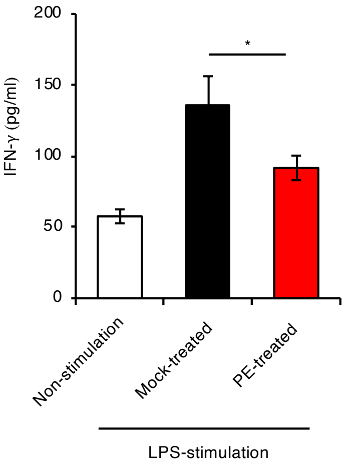

**Supplemental Figure 4**. **PE has immunomodulatory effects against mouse sepsis model.**

Blood serum were collected from Mock-treated mice (n=2), LPS-treated mice (n=3) or LPS and PE-treated mice (n=3). IFN-γ levels in the serum were assessed by use of an ELISA. * $p < 0.05$; (Mann-Whitney U test).
